# Proteome integral solubility alteration assay combined with multi-criteria decision-making analysis for developing adverse outcome pathways

**DOI:** 10.1101/2022.10.17.512512

**Authors:** Veronica Lizano-Fallas, Ana Carrasco del Amor, Susana Cristobal

**Affiliations:** Department of Biomedical and Clinical Sciences, Cell Biology, Faculty of Medicine, Linköping University, Linköping, Sweden; Ikerbasque, Basque Foundation for Sciences, Department of Physiology, Faculty of Medicine, and Nursing, University of the Basque Country UPV/EHU, Spain

**Keywords:** Proteome integral solubility alteration assay, analytic hierarchy process, adverse outcome pathways, TCDD, thermal proteome profiling, risk assessment

## Abstract

Understanding the biological impact of chemicals is hindered by the high number and diversity of compounds in the market. To simplify the chemical risk assessment, the adverse outcome pathway (AOP) method has arisen as a framework to predict the impact of chemical exposure on human and environmental health. The development of this predictive tool requires knowledge of the molecular interaction between chemicals and protein targets. Those molecular initiating events connect alterations of cellular function with physiological impairment. This strategy aims to focus on the complex biological interaction to predict the impact on health. The high-throughput identification of all chemical targets can be obtained by a proteomics-based thermal shift assay, however, selecting the priority target candidate is a biased process strongly dependent on expert knowledge and literature. Here, we unravel new molecular initiating event from a tested chemical combining the target deconvolution by the proteome integral solubility alteration (PISA) assay, and the target selection by an analytical hierarchy process (AHP) approach. In the proof-of-concept study, we identified by PISA assay 8 protein targets for 2,3,7,8 tetrachlorodibenzo-p-dioxin (TCDD) from the soluble proteome from hepatic cells containing 2824 proteins. The definition of the AHP approach facilitates the selection of heat shock protein beta-1 (Hspb1) as the most suitable protein for developing AOPs. Our results demonstrated that the process of target identification is independent from a chemical characterization, and that the process of data curation and target selection is less sensitive to lack of toxicological information. We anticipate that this innovative integration of methods could decipher the chemical-protein interactions from new chemicals including the new alternative chemicals designed for chemical replacement and that would discover new molecular initiating events to support more sustainable methodologies to gain time and resources in chemicals assessment.

**SYNOPSIS:** Our combined methodologies can determine the most suitable target to develop adverse outcome pathways from the proteome-wide protein target identification.

## 1. INTRODUCTION

Chemicals are widely used and offer significant benefits to our daily lives. However, the number, diversity and complexity of substances coming to the market is increasing enormously. Only under the European Union legislative frameworks, more than 200 000 chemicals are registered ^1^. Consequently, the concern about human and environment safety is rising. The risk to human health and the environment of many of these compounds is still unknown and their assessment through traditional animal testing approaches is not practical in terms of time- and cost-consuming and does not support the 3Rs principles of the use of laboratory animals: replacement, reduction, and refinement ^2^. Moreover, the thousands of new and existing chemicals that required to be evaluated, the financial support assigned for this kind of assessment is not increasing ^3^. Thus, predictive toxicology based on mechanistic information has become critical ^4^.

Under this context, the adverse outcome pathway (AOP) framework represents an approach that fulfills the mentioned requirements, and it is expected to support in the deconvolution of the twenty-first century challenges regarding chemical risk assessment ^2,5^. The AOP concept was introduce in 2010 in the field of ecotoxicology and relies on existing knowledge to provide a framework to compile, organize and assess significant information on biological and toxicological effects of chemicals. The generic structure of an AOP portrays the linkage between a molecular initiating event (MIE) and an adverse outcome at a biological level of organization relevant to risk assessment, i.e., at the level of organism or population, passing across key events and key event relationships ^6,7^.

Due to the multiple applications of AOPs in toxicology, ecotoxicology, and risk assessment its use has increased and gained widespread acceptance. In the last 5 years, the number of AOPs has been doubled reaching 407, and the number of stressors associated to trigger a MIE has been increased from 250 to 741 ^4,8^. Such associations or MIEs characterization rely on *in silico, in chemico*, and *in vitro* data ^4,9^ and over the past decades, significant advances have been done in the development of these type of methodologies. One of these advances includes the conception of the term high-throughput toxicology. It describes the application of batteries of *in vitro* bioassays, using high throughput technologies, to scrutinize rapidly and cost-effectively the interaction of individual chemicals with specific molecular targets or biological pathways whose perturbation could lead to adverse outcomes ^3,10^.

Under this context, the programs ToxCast and Tox21 were developed to evaluate chemicals, using batteries of around 700 and 100 *in vitro* bioassays for specific molecular targets or biological pathways, respectively ^10,11^. More than 10,000 individual chemicals have been assessed since 2008 within these programs after the shift from traditional toxicology to high-throughput toxicology. These promising achievements lead to believe that most of individual chemicals in the market could be screened for a broad spectrum of toxicologically relevant biological activities by 2026 ^10^. However, evaluating more than 100,000 chemicals was a task implausible to achieve ^11^.

Maintaining the approach of high-throughput toxicology, a possible solution to this challenging task could be moving a step forward to the use of omics techniques. One of the advantages of using omics instead of large suites of in vitro bioassays is that with omics only one single analytical assay is performed, and all biological pathways are covered at the same time, gaining in time, resources, and completeness of the acquired knowledge. An example of an omic methodology recently utilized to screen chemicals is the proteome integral solubility alteration (PISA) assay ^12^.

This approach is based on the application of the thermal shift principles at a proteome scale for identifying protein targets of bioactive compounds. Proteins direct biological processes through the interaction with small molecules in the cell ^13^. Therefore, chemicals entering the cellular environment could interact with proteins, altering their solubility and consequently, their function.

This solubility alteration can be probed by applying a stability factor, as temperature in thermal profiling ^14^. Upon this principle, in this previous study, PISA was applied for the first time in the field of toxicology and showed its potency by predicting the impact of exposure to individual chemicals, mixtures of chemicals, new chemicals from biodiscovery, and side-effects of newly developed drugs ^12^. Even though the PISA methodology offers completeness by assessing the entire proteome, covering all biological pathways, an approach that could extract the uniqueness out of the complete answer is required for transferring and integrate this knowledge into AOPs by the identification of MIEs.

An approach that could assist the selection of an individual MIE or the most valuable MIEs from the high-throughput data considering relevant aspects for developing an AOP could be a potential solution. Systematic quantitative methods to select or rank alternatives among several options contemplating different factors are known as multi-criteria decision-making analysis (MCDM) techniques. Any problem where a significant decision is required can be solved by the application of MCDM methods ^15^. According to Roy, decisions can be classified in four main types: choice problem, sorting problem, ranking problem, and description problem. In the case of choice problems, the aim is to select the single best option or reduce the group of alternatives to a subset of equivalent good options. Within sorting problems, the alternatives are grouped into categories. In ranking problems, the options are ordered from best to worst by means of scores or pairwise comparisons. While in description problems, the aim is to describe the alternatives and their consequences ^16^.

Based on this classification, many MCDM methods have been developed. Aspects to consider for choosing the appropriate methodology are the type of problem to be solved, the desired output quality and the modelling effort able to input. The analytic hierarchy process (AHP) belongs to a family of decision-making tools for choice and ranking problems and offers the highest output quality requiring a medium modelling effort to perform the analysis, making it suitable for assist the selection of a MIE from the high-throughput data. Within AHP the best alternative is selected by enumerating key factors for decision making and assessing the relative value of different decision alternatives, integrating evidence-based data ^15,17^. Previously, MCDM techniques have been used for the toxicity prioritization of fine dust (PM2.5) sources, ranking chemicals for toxicological impact assessments, and assessing the risk from multi-ingredient dietary supplements, but to the knowledge of the authors, it has never been applied to the field of AOPs before ^18–20^.

In the present study, we propose to accelerate and improve chemicals assessment by the combination of a high-throughput based-proteomics method for target deconvolution, the PISA assay, and a multiple-criteria decision-making analysis technique, AHP, to unravel new MIEs which could be used to develop new AOPs.

We anticipate that this innovative combination of techniques would help to develop new AOPs and fill out missing information of the preexisting ones, benefiting the potential applications of the AOP framework in chemical risk assessment, such as chemical grouping to predict the toxicological properties of a target substance; chemical prioritization to undergo more detailed testing; safety assessment predictions by the generation of integrated approaches to testing and assessment; development or refinement of methods to test specific key events; development of alternatives to animal testing; among others ^2,4^.

## 2. MATERIALS AND METHODS

### 2.1. Sample preparation

Reagents and medium were purchased from Sigma-Aldrich (Sant Louis, MO, USA), unless otherwise noted. PBS was purchased from Trevigen (Gaithersburg, MD, USA). HepG2 cells were grown until 70% confluence in EMEM medium supplemented with 7% fetal bovine serum (ATTC), 1675 mM L-glutamine, 85 U/mL penicillin and 85 μg/mL streptomycin. Cells were harvested and centrifugated at 340 g for 4 min at 4 °C. Three washes were made with 30 mL cold PBS. Resuspension and centrifugation at 340 g for 4 min at 4 °C. Washed pellets were snap frozen in liquid nitrogen and stored at −80 °C until lysis.

### 2.2. Selection of test compound and concentration

The bioactive compound analysed in this study was 2,3,7,8 tetrachlorodibenzo-p-dioxin (TCDD), a persistent organic pollutant and endocrine disruptor compound and the corresponding highest concentration tested was 25 nM.

The rationale for the selection of the test compound was to include a well-known xenobiotic of high relevance for human and environmental health. Therefore, we included TCDD and its concentration from a previous study for *in vitro* exposures to human hepatic cell line HepRG. Concentration selection was based on the translation of external intakes into internal doses in hepatic cells ^21^.

### 2.3. Two Dimensions Proteome Integral Solubility Alteration (2D PISA) experiments in HepG2 cells protein extracts

HepG2 cells were resuspended in ice-cold PBS and lysed in ice bath by sonication in cycles of 10 s on /5 s off for 1 min at 6–10 μm amplitude at 25 % intensity from an exponential ultrasonic horn of 3 mm in a Soniprep 150 MSE (MSE Ltd., Lower Sydenham, London, UK). The insoluble parts were sedimented by centrifugation at 100,000 g for 60 min at 4 °C 22. Protein concentration was determined by BCA assay ^23^. The soluble proteome was used to perform the 2D PISA assay, as described in Gaetani *et al.* ^14^ with some modifications. Briefly, the soluble proteome and the studied chemical were incubated for 10 min at 25 °C. The incubation was performed with TCDD at 10 different concentrations. The studied concentrations were 100, 90, 80, 70, 60, 50, 40, 30, 20 and 0 %. The highest concentration (100 %) corresponds to 25 nM TCDD. The control sample (0%) was incubated in the presence of the vehicle, dimethyl sulfoxide (DMSO), utilized for the solubilization of the compound. Ten specific temperatures were selected for the thermal assay: 37, 42, 46, 49, 51, 53, 55, 58, 62 and 67 °C. These temperatures were selected to ensure that at least 90% of the studied proteins have their melting point within this range ^24^. Aliquots containing 10 μg of protein (one for each of the temperatures in the entire range covered in the thermal shift assay) were independently heated at the corresponding temperature for 3 min, followed by 3 min at room temperature. For each concentration, aliquots of all temperature points were pooled and centrifuged at 100,000 g for 20 min at 4 °C, to remove the proteins that had an alteration in solubility after the thermal shift assay ^24^. Supernatants from intermediate concentrations were combined. The collection of soluble fractions in the supernatants from the three conditions (control – 0 %, intermediate concentrations and highest concentration – 100 %) were processed using a general bottom-up proteomics workflow and the purified peptides were analyzed by label-free nano liquid chromatography-tandem mass spectrometry analysis (nLC-MS/MS). Three biological replicates were performed for each experiment ^25^.

### 2.4. Filter aided sample preparation (FASP)

The samples were digested following the FASP method. First, the protein samples corresponding to the supernatants after centrifugation were prepared with SDT buffer (2% SDS, 100 mM Tris-HCl, pH 7.6 and 100 mM DTT), according to Wiśniewski *et al.* ^26^. To perform FASP, the samples were diluted with 200 μl of 8 M urea in 0.1 M Tris/ HCl, pH 8.5 (UA) in 30 kDa microcon centrifugal filter units. The filter units were centrifuged at 14,000 g for 15 min at 20 °C. The concentrated samples were diluted with 200 μl of UA and centrifuged at 14,000 g for 15 min at 20 °C. After discharging the flow-through 100 μl of 0.05 M iodoacetamide was added to the filter units, mixed for 1 min at 600 rpm on a thermo-mixer, and incubated static for 20 min in dark. The solution was drained by spinning the filter units at 14,000 g for 10 min. The filter units were washed three times with 100 μl buffer UA and centrifuged at 14,000 g for 15 min. The filter units were washed three times with 100 μl of 50 mM ammonium bicarbonate. Endopeptidase trypsin solution in the ratio 1:100 was prepared with 50 mM ammonium bicarbonate, dispensed, and mixed at 600 rpm in the thermomixer for 1 min. These units were then incubated in a wet chamber at 37 °C for about 16 h to achieve effective trypsinization. After 16 h of incubation, the filter units were transferred into new collection tubes. To recover the digested peptides, the tubes were centrifuged at 14,000 g for 10 min. Peptide recovery was completed by rinsing the filters with 50 μl of 0.5 M NaCl and collected by centrifugation. The samples were acidified with 10% formic acid (FA) to achieve pH between 3 and 2. The desalting process was performed by reverse phase chromatography in C18 top tips using acetonitrile (ACN; 60% v/v) with FA (0.1% v/v) for elution, and vacuum dried to be stored at −80 °C till further analysis.

### 2.5. Nano LC-MS/MS analysis

The desalted peptides were reconstituted with 0.1 % FA in ultra-pure milli-Q water and the concentration was measured using a Nanodrop (Thermo Scientific). Peptides were analyzed in a QExactive quadrupole-orbitrap mass spectrometer (Thermo Scientific). Samples were separated using an EASY nLC 1200 system (Thermo Scientific) and tryptic peptides were injected into a pre-column (Acclaim PepMap 100 Å, 75 um × 2 cm) and peptide separation was performed using an EASY-Spray C18 reversed-phase nano LC column (PepMap RSLC C18, 2 um, 100 Å, 75 um × 25 cm). A linear gradient of 6 to 40% buffer B (0.1 % FA in ACN) against buffer A (0.1 % FA in water) during 78 min and 100% buffer B against buffer A till 100 min, was carried out with a constant flow rate of 300 nl/min. Full scan MS spectra were recorded in the positive mode electrospray ionization with an ion spray voltage power frequency (pf) of 1.9 kV (kV), a radio frequency lens voltage of 60 and a capillary temperature of 275 °C, at a resolution of 30,000 and top 15 intense ions were selected for MS/MS under an isolation width of 1.2 m/z units. The MS/MS scans with higher energy collision dissociation fragmentation at normalized collision energy of 27 % to fragment the ions in the collision induced dissociation mode.

### 2.6. Peptide and protein identification and quantification

Proteome Discoverer (v2.1, Thermo Fischer Scientific) was used for protein identification and quantification. The MS/MS spectra (. raw files) were searched by Sequest HT against the Human database from Uniprot (UP000005640; 95,959 entries). A maximum of 2 tryptic cleavages were allowed, the precursor and fragment mass tolerance were 10 ppm and 0.6 Da, respectively. Peptides with a false discovery rate (FDR) of less than 0.01 and validation based on q-value were used as identified. The minimum peptide length considered was 6 and the FDR was set to 0.1. Proteins were quantified using the average of top three peptide MS1-areas, yielding raw protein abundances. Common contaminants like human keratin and bovine trypsin were also included in the database during the searches for minimizing false identifications. The mass spectrometry proteomics data have been deposited to the ProteomeXchange Consortium via the PRIDE ^27^ partner repository with the dataset identifier PXD033056.

### 2.7. Analysis of 2D PISA assay

Two dimensions PISA assay measures the protein abundance from 3 biological replicates of 3 conditions (control – 0 %, intermediate concentrations and highest concentration – 100 % of the tested compound). Protein abundances from control and the highest concentration represent, for each protein, the integral of the area under its melting curve within the used temperature interval. If S_m_ is the value for the control condition and S_m_’ is the corresponding value for the highest concentration condition, then the PISA analogue of the melting temperature shift (ΔT_m_) is

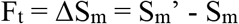

Protein abundance from intermediate concentrations (S_m_”) represents an integral of the concentration-dependance curve. Similarly, the PISA analogue of the compound concentration required to induce half of the ΔT_m_ (C0) is

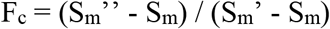

For each protein, the abundance was normalized on the average value for the control condition, and then F_t_ and F_c_ were calculated as described. Two-tailed Student’s t-test (with equal or unequal variance depending on F-test) was applied to calculate p-values. Proteins with p-values < 0.05 for both F_t_ and F_c_ were considered protein targets, meaning to be the proteins combining solubility alteration with action at a low compound concentration ^14^. The data was represented in a scatter plot combining Ft and Fc p-values, to facilitate the visualization of the protein targets.

### 2.8. Nanoscale Differential Scanning Fluorometry (NanoDSF) protein-chemical binding validation

One identified protein target was selected to perform the protein-chemical binding validation. We assessed differential scanning fluorimetry with a nanoDSF device as an orthogonal validation approach. NanoDSF is based on the changes in the intrinsic tryptophan fluorescence (ITF) resulting from alterations of the 3D-structure of proteins, when proteins unfold, as a function of the temperature. Therefore, a melting temperature (Tm) can be determined ^28^.

Monitoring of the ITF at 330 nm and 350 nm during protein thermal denaturation was carried out in a Prometheus NT.48 instrument from NanoTemper Technologies with an excitation wavelength of 280 nm. Excitation power was set at 25 %. Capillaries were filled with 10 μl of a solution containing the protein and TCDD, placed into the sample holder and a temperature gradient of 0.5 °C/min from 20 °C to 80 °C was applied. The ratio of the recorded emission intensities (Em350nm/Em330nm), which represents the change in tryptophan fluorescence intensity was plotted as a function of the temperature. The fluorescence intensity ratio and its first derivative were calculated with the manufacturer’s software (PR.ThermControl, version 2.3.1). For validating protein-chemical binding, purified protein was mixed with 3 different concentrations of TCDD. The tested compound concentrations were 5 nM (20 %), 15 nM (60 %), and 25 nM (100 %). Protein final concentration was 0.5 mg/ml. Control was performed with purified protein in PBS and DMSO maintaining the corresponding protein concentration as for the TCDD. Three replicates were carried out for each condition.

Selection of the protein target for validation was based on the availability on the market (full-length recombinant protein and without GST tag, due to possible interferences with chemical binding) and the number of tryptophan residues (at least 2).

### 2.9. Selection of protein target for new Adverse Outcome Pathways

From the identified protein targets for TCDD, one was selected as the best protein for developing an AOP by the multiple-criteria decision-making analysis technique, AHP. For a better understanding of the methods, the workflow is shown in Figure 1.

**Figure 1.**
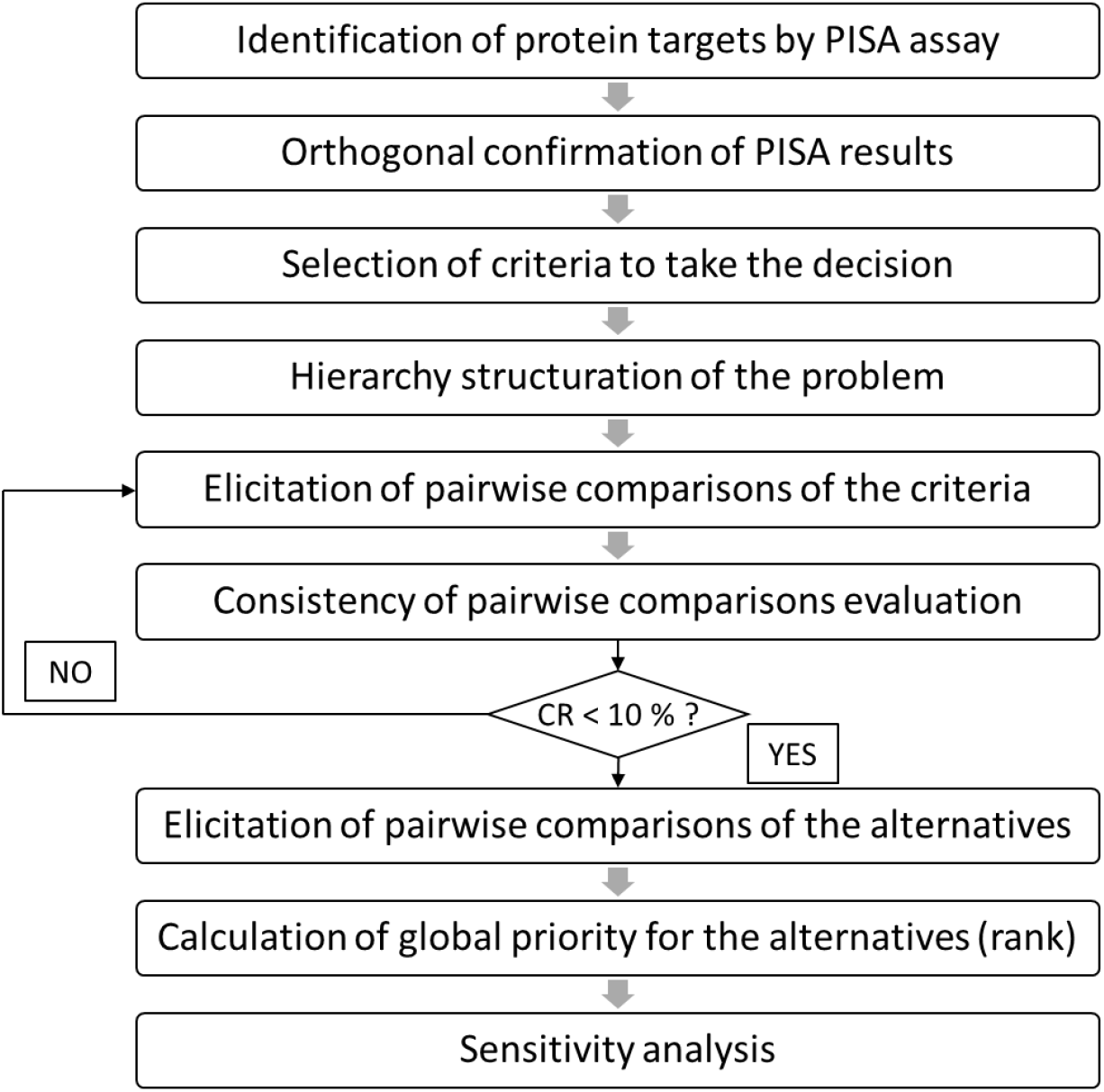
Workflow followed to select from identified protein targets by 2D PISA assay the best one for developing AOPs. CR: consistency ratio. Adapted from Yadav and collaborators ^31^.

The AHP approach arrange the factors considered to take a decision in a hierarchic structure and relies on three steps ^17^. The first one is the decomposition. Here, the problem was structured as a hierarchy, where the first level contains the overall goal, i.e., selection of the best protein for developing an AOP. The following level corresponds to the criteria which contribute to the goal. These criteria were chosen by the authors based on the requirements and guidelines for developing an AOP ^4,9,29^. And the third level includes the alternatives (protein targets) to be evaluated in terms of the criteria in the second level.

The second step is the elicitation of pairwise comparison judgments, where a matrix of the relative importance of each criterion over each other was performed using a scale from 1 to 9, according to the expertise of the authors. In this scale, 1 denotes that the two factors contribute equally to the goal, 9 represents extreme importance of one over another, while 3 indicates slight importance. A numeric scale of 5 represents moderate importance and 7 indicates a very strong relevance of one factor over another. The values 2, 4, 6 and 8 represent the intermediate values between two adjacent judgments.

After calculating the priority vector of the matrix, the consistency of the pairwise comparisons was evaluated through the calculation of the consistency ratio (CR), which involved the following operations:

- Computing the principal eigenvalue (λmax) as in eq 1.

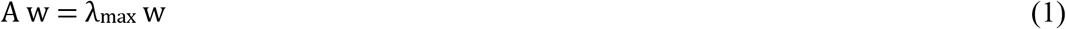 Where *A* is the priority vector, and *w* are the eigenvalues of the vector *A*
- Computing the consistency index (CI) as in eq 2

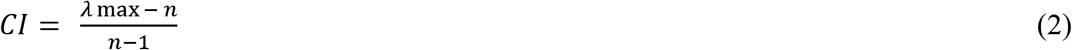 Where *n* is the number of criteria
- Calculation of the CR as in eq 3

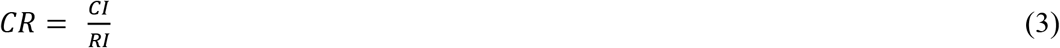

Where *RI* corresponds to the appropriate value of the random consistency indices i.e., the CI expected from a matrix of that order. According to Saaty the value of *RI* is 0 up to order 2 while for 3 to 10 the random consistency index values are 0.58, 0.90, 1.12, 1.24, 1.32, 1.41, 1.45 and 1.49, respectively. A consistency ratio up to 10 % is considered acceptable ^30^.

The third step of AHP is to establish the global priorities of the alternatives. This was done by laying out the local priorities of the alternatives with respect to each criterion in a matrix (by pairwise comparison judgments using a scale from 1 to 9). Then, the local priorities are multiplying by the priority of the corresponding criterion and adding across each row. The alternative with the highest global priority was selected as the best protein for developing an AOP.

Finally, to increase the viability and robustness of the results, a sensitivity analysis was performed. The sensitivity analysis assesses the effects on the final decision after the minor variation in the input. Any slightest change in the current priority can alter the existing global priorities of the alternatives ^31^. Here, the criterion with the highest priority was selected and varied from 0.05 to 0.9 in intervals of 0.05 to calculate the global priorities of the alternatives for each interval. If the alternative selected as the best protein for developing an AOP maintains its position at every interval the result of the AHP method is validated.

## 3. RESULTS AND DISCUSSION

### 3.1. *In chemico* protein targets of TCDD in the soluble proteome from hepatocytes

We evaluated the 2D PISA assay using temperature as stability factor to identify the proteins from the HepG2 cell line soluble proteome that interact with TCDD. The compound was evaluated on 10 different concentrations, in a dilution series of 10%, starting from 20% of the highest concentration - 25 nM (100%). We used three biological replicates per condition, and the corresponding control with a vehicle.

The soluble proteome analyzed yielded 2,824 proteins identified and quantified across the replicates, and 1,475 proteins quantified in all three replicates were included in the 2D PISA analysis. After the analysis, 8 proteins were identified as targets for TCDD. The protein targets can be depicted from the plot (Figure 2). The protein target general transcription factor 3C polypeptide 4 (Gtf3c4) showed the highest solubility alteration at the highest concentration tested of TCDD and an action at lower concentrations of TCDD. The other protein targets are heat shock protein beta-1 (Hspb1), ras-related protein Rab-1A (Rab1a), proteasome adapter and scaffold protein ECM29 (Kiaa0368), myotrophin (Mtpn), protein FAM98B (Fam98b), glyoxylate reductase/hydroxypyruvate reductase (Grhpr), protein canopy homolog 3 (Cnpy3).

**Figure 2.**
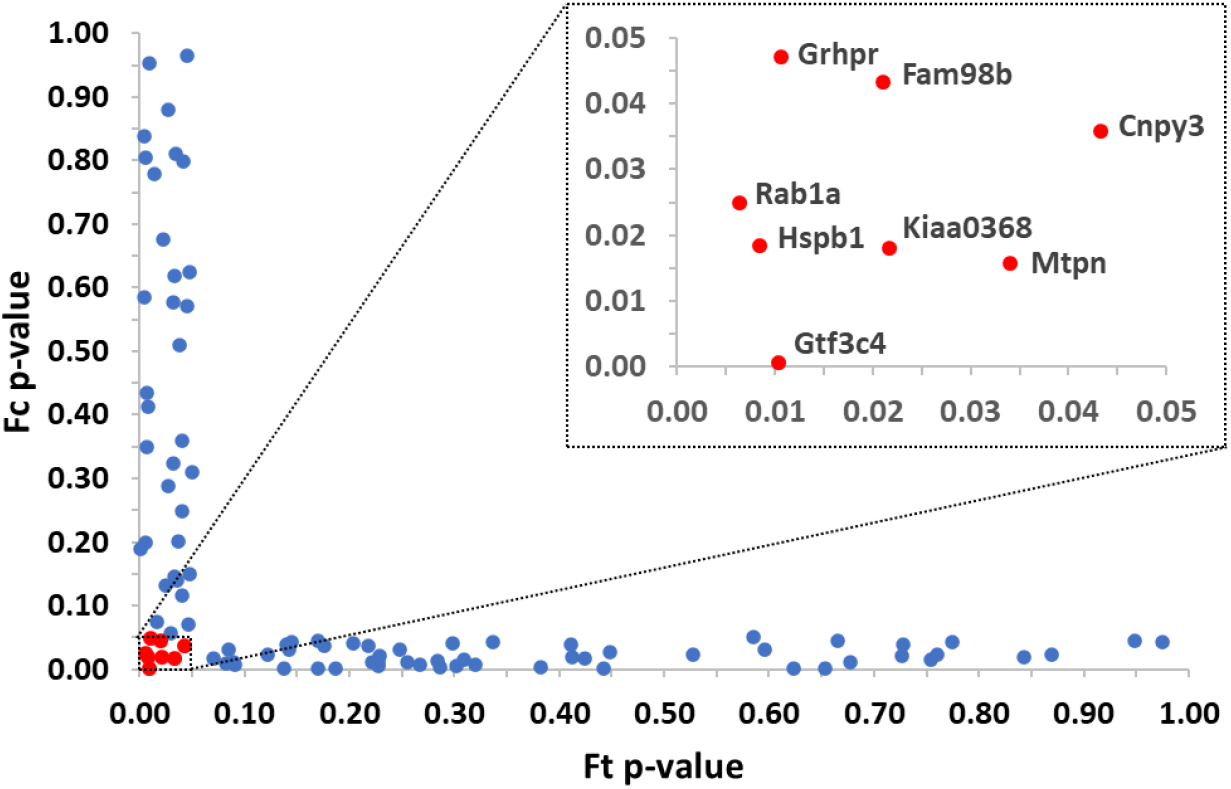
Protein targets identified by 2D PISA method for TCDD and the soluble proteome from hepatocytes. The studied concentrations were 100, 90, 80, 70, 60, 50, 40, 30, 20 and 0 %. The highest concentration (100 %) corresponds to 25 nM TCDD. Identified protein targets shown in red and labeled. (For interpretation of the references to color in this figure legend, the reader is referred to the web version of this article.)

TCDD is classified as a persistent organic pollutant and endocrine disruptor compound and offers a well-known AOP based on the binding and consequently activation of the aryl hydrocarbon receptor (AOP 21) ^32^, is therefore expected to get this protein as target. However, in our study, this protein cannot be obtained as one of the targets because this specific protein is not part of the studied proteome. Although PISA method is a holistic and unbiased approach to identify protein targets, the results are constraint to the size of the studied proteome. In this study, we analyzed a proteome composed of 2,824 proteins for its interactions with TCDD with our methodology. The introduction of a pre-fractionation prior to LC-MS/MS analysis would have enlarged the proteome size ^14^ but also increased time and cost, compromising the method applicability to large screening of chemicals. The application of PISA to toxicology requires a tight balance between what is the level of completeness of the cellular proteome that we scrutinize in a single experiment without reducing the specificity and sensitivity of the method. We have previously reported that the identification of targets from hydrophobic chemicals increased in robustness by introducing a step of sedimentation of the microsomal vesicles before the thermal shift assay. This implementation of the method eliminates the risk of the unquantified hydrophobic chemical sequestration inside the microsomes, that is a temperature dependent process, and compromised the maintenance of the stable concentration of chemical and proteome along the thermal shift assay ^12,22^. Therefore, our PISA method has been implemented for its application to toxicology to obtain a soluble proteome that would be stable with chemical compounds and without larger pre-fractionation that would increase costs. Our results showed unbiased identification of any interaction between TCDD and all the proteins in the studied proteome, giving us the chance to unravel novel molecular initiating events, that would have never been studied with a single protein assay.

### 3.2. Orthogonal protein-chemical binding validation with NanoDSF

Protein melting temperature modification is expected when protein stability changes because of protein-chemical interactions. Therefore, nanoDSF can be used as a protein-chemical binding validation approach. For this purpose, from the 2D PISA assay, the protein target heat shock protein beta-1 (Hspb1) for TCDD was selected to perform the binding validation with NanoDSF.

Hspb1 have 6 tryptophan residues, condition required for this analysis. Purified protein was purchased from Novus Biological as an un-tagged full-length recombinant protein (NBC1-18364). After performing nanoDSF validation, compared to the controls, a shift in the melting temperature was observed from the first derivative of the ratio of the emission intensities (Em350nm/Em330nm) of the interaction between Hspb1 and TCDD at 3 different concentrations, as expected (Figure 3). The classical process for individual target validation involves determining target engagement in *in vitro* or *in vivo.* Here, the purpose of utilizing an orthogonal method is confirming the chemical and protein interaction at the structural level. However, this is not an alternative for target validation as it did not include analysis of effects of the interaction. It is a confirmation of the results from PISA method that are constraint to chemical-protein interactions ^14^. The results from our methodology are presented as a list of proteins and not just a single unique target. Many of them are proteins never described as targets in the literature. Therefore, attempts to validate the results should imply different strategies. Our focus here is improving our mechanistic understanding of the MIE through target identification. Therefore, the next step is analyzing the PISA results with MCDM techniques that facilitates the discovering of the most suitable target to feed the AOP framework.

**Figure 3.**
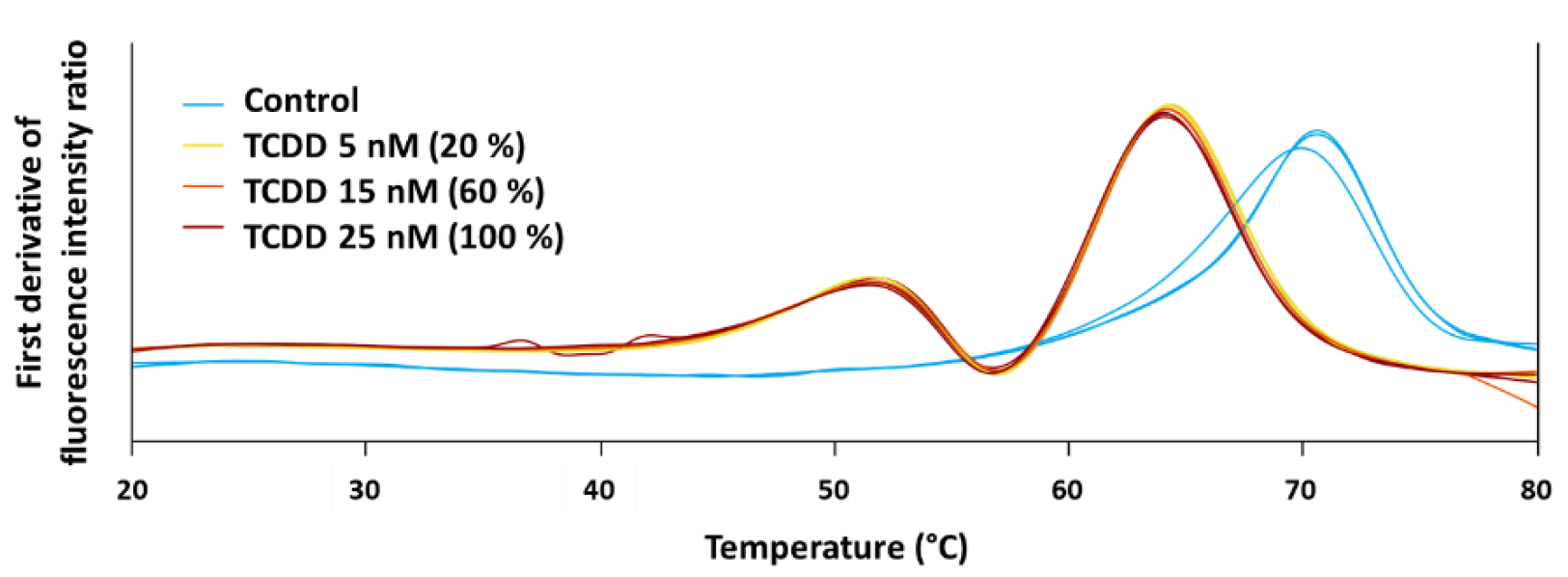
Melting temperatures obtained from nanoDSF protein-chemical binding validation for the interaction of Hspb1 with TCDD at 5 nM (20 %) (yellow), 15 nM (60 %) (orange), and 25 nM (100 %) (red). A control without TCDD was included (light blue). (For interpretation of the references to color in this figure legend, the reader is referred to the web version of this article.)

### 3.3. Selection of the best protein target for developing AOPs

From the 8 identified protein targets for TCDD by 2D PISA assay, one was selected by the AHP method as the best protein for the development of an AOP. The first challenge for the application of AHP was to generate a hierarchical structure of the problem. The first level of the hierarchy was the overall goal of the analysis, i.e., selecting the best protein target to develop an AOP. The second level contained 8 criteria that were selected based on the requirements and guidelines for developing an AOP ^4,9,29^. These criteria are described in Table 1. The third level included the 8 protein targets identified by the 2D PISA assay as the alternatives to be evaluated in terms of the criteria of the second level.

**Table 1.** Description of the selected criteria which contribute to the overall goal of selecting by AHP from identified protein targets by 2D PISA assay the best one for developing AOPs.

| Criteria | Source | Description |
| --- | --- | --- |
| Position in Ft (solubility alteration) ranking | 2D PISA assay | Protein with highest solubility alteration (lowest position in ranking) is more likely to bind the chemical |
| Position in Fc p-value ranking | 2D PISA assay | Protein with the highest significance (lowest position in ranking) for solubility alteration is more likely to bind the chemical |
| Number of diseases where it is involved | UniProt | Protein with the highest number of diseases where it is involved has more relevance to be included in an AOP |
| Number of reported negative effects on cells/organs/organisms when its functionality is absent | Literature | Protein with the highest number of reported negative effects on PubMed under the search criteria "lack/absence/knockdown/depletion/knockout of name of the protein" has more relevance to be included in an AOP |
| Relevance of reported negative effects on cells/organs/organisms when its functionality is absent | Expertise criteria | Negative effects with regulatory significance (accepted protection goal or common apical endpoint in an established regulatory guideline study) are more relevant |
| Number of pathways where it has participation | Reactome / Metabolic Atlas | Protein with the highest number of pathways where it has participation has more relevance to be included in an AOP |
| Relevance of pathways where it has participation | Expertise criteria | Pathways associated with adverse outcomes are more relevant |
| Number of functional and physical protein associations with other protein targets | STRING | Protein with highest number of protein associations with other protein targets has more relevance to be included in an AOP |

After defining the 8 criteria for selecting the best protein target for developing an AOP, a matrix of pairwise comparison judgments of the criteria was performed by the authors expertise. Table 2 shows the obtained matrix, the priority vector and the corresponding λ_max_, CI, RI, and CR. According to the priority vector, the criteria: position in Ft (solubility alteration) ranking and position in Fc p-value ranking obtained the highest weight of relevance for a protein to be used for developing AOPs. Those high weight values relate to the level of importance of evidence that protein-chemical interaction data could provide for a MIE. The molecular confirmation of the MIE by PISA here is therefore key in a bottom-up strategy that starts from the molecular event. However, top-down strategies starting from an observed adverse outcome has been more frequently applied to AOP development 9 based on the difficulties in isolate and identify molecular interactions ^29^.

**Table 2.** Pairwise comparison matrix for the criteria at level 2 of the hierarchy and the computed values of priority vector, λ_max_, CI, RI, and CR.

|  | Position in Ft (solubility alteration) ranking | Position in Fc p-value ranking | Number of diseases where it is involved | Number of reported negative effects on cells/organs/organisms when functionality is absent | Relevance of reported negative effects on cells/organs/organisms when functionality is absent | Number of pathways where it has participation | Relevance of pathways where it has participation | Number of functional and physical protein associations with other protein targets | Priority vector |
| --- | --- | --- | --- | --- | --- | --- | --- | --- | --- |
| Position in Ft (solubility alteration) ranking | 1 | 1 | 5 | 5 | 3 | 5 | 3 | 7 | 0.258 |
| Position in Fc p-value ranking | 1 | 1 | 5 | 5 | 3 | 5 | 3 | 7 | 0.258 |
| Number of diseases where it is involved | 1/5 | 1/5 | 1 | 1 | 1/3 | 2 | 1/3 | 5 | 0.060 |
| Number of reported negative effects on cells/organs/organisms when functionality is absent | 1/5 | 1/5 | 1 | 1 | 1/5 | 3 | 1/5 | 5 | 0.061 |
| Relevance of reported negative effects on cells/organs/organisms when functionality is absent | 1/3 | 1/3 | 3 | 5 | 1 | 5 | 3 | 7 | 0.166 |
| Number of pathways where it has participation | 1/5 | 1/5 | 1/2 | 1/3 | 1/5 | 1 | 1/5 | 5 | 0.044 |
| Relevance of pathways where it has participation | 1/3 | 1/3 | 3 | 5 | 1/3 | 5 | 1 | 7 | 0.133 |
| Number of functional and physical protein associations with other protein targets | 1/7 | 1/7 | 1/5 | 1/5 | 1/7 | 1/5 | 1/7 | 1 | 0.021 |
| $\lambda_{\max} = 8.831$ CI = 0.119 RI = 1.41 CR = 0.084 | | | | | | | | | |

The following criteria in the priority rank were: i) relevance of reported negative effects on cells/organs/organisms when protein functionality is absent, ii) relevance of pathways where the protein has participation, iii) number of reported negative effects on cells/organs/organisms when protein functionality is absent, and iv) number of diseases where the protein is involved. This order in the priority rank is associated to how an observed adverse outcome of a chemical is relevant from a risk assessment perspective i.e., it corresponds to an accepted protection goal or common apical endpoint in an established regulatory guideline study ^4,29^. Finally, the lowest weight was occupied by the criterion number of functional and physical protein associations with other protein targets. This is quantitative data available to retrieved for most of the proteins. This information could be of useful in the implementation of AOP networks, is not directly required to the AOP development.

The third step of AHP is to establish the global priorities of the alternatives by pairwise comparison judgments. *A priori*, a database containing the information of each protein for each criterion was created to perform the comparisons (Supplementary Table 1). The main sources utilized to retrieve the information for the database are indicated in Table 1. The matrices of pairwise comparison judgments of the alternatives were performed by the authors, and the corresponding local priority vectors, λ_max_, CI, RI, and CR are shown in Supplementary Table 2. The local priorities were multiplying by the priority of each criterion. The obtained values were added to derive the global priority of each protein (Table 3). Protein Hspb1 reached the highest global priority (0.414), turning it into the alternative selected by the AHP strategy as the best protein for developing an AOP. However, if the selection would have been based on the 2D PISA assay ranks that are only based on solubility alteration and degree of statistical significance, this target would not occupy the first position. This result shows the relevance of using an integral, systematic approach, where other aspects of the protein further than chemical binding are included. Hspb1 reported 10 negative effects on cells/organs when its functionality is absent, most of them with regulatory significance. Other alternatives reported maximum 2 negative effects. Furthermore, it is associated with 4 pathways while other proteins are associated with maximum 1, except from protein Rab1a which also is related to 4 pathways. Protein Hspb1 has no functional and physical protein associations with other protein targets, however its position in the global priority rank of AHP is not affected since this criterion has the lowest weight.

**Table 3.** Local priority for each criterion and global priority of the protein targets (alternatives). Criteria priority vector shown bold.

| Protein | Local priority |  |  |  |  |  |  |  | Global priority |
| --- | --- | --- | --- | --- | --- | --- | --- | --- | --- |
|  | Position in Ft (solubility alteration) ranking | Position in Fc p-value ranking | Number of diseases where it is involved | Number of reported negative effects on cells/organs/organisms when functionality is absent | Relevance of reported negative effects on cells/organs/organisms when functionality is absent | Number of pathways where it has participation | Relevance of pathways where it has participation | Number of functional and physical protein associations with other protein targets |  |
|  | **0.258** | **0.258** | **0.060** | **0.061** | **0.166** | **0.044** | **0.133** | **0.021** |  |
| Gtf3c4 | 0.093 | 0.130 | 0.006 | 0.006 | 0.016 | 0.004 | 0.008 | 0.002 | 0.265 |
| Fam98b | 0.067 | 0.019 | 0.006 | 0.006 | 0.016 | 0.004 | 0.015 | 0.002 | 0.135 |
| Hspb1 | 0.048 | 0.064 | 0.058 | 0.052 | 0.118 | 0.035 | 0.037 | 0.002 | 0.414 |
| Rab1a | 0.035 | 0.047 | 0.006 | 0.022 | 0.062 | 0.035 | 0.037 | 0.019 | 0.264 |
| Mtpn | 0.024 | 0.103 | 0.006 | 0.006 | 0.016 | 0.001 | 0.002 | 0.010 | 0.168 |
| Kiaa0368 | 0.016 | 0.082 | 0.006 | 0.011 | 0.016 | 0.000 | 0.001 | 0.010 | 0.142 |
| Grhpr | 0.010 | 0.011 | 0.042 | 0.018 | 0.149 | 0.004 | 0.055 | 0.002 | 0.291 |
| Cnpy3 | 0.007 | 0.035 | 0.042 | 0.018 | 0.149 | 0.004 | 0.041 | 0.002 | 0.297 |

Besides, AHP results were validated through a sensitivity analysis. Due to the criterion position in Ft (solubility alteration) ranking had the maximum priority it was used as input to assess the minor variation on the final decision, varying it from 0.05 to 0.9 in intervals of 0.05 to calculate the global priorities of the alternatives for each interval. Table 4 shows that a minor variation occurs in the ranking of proteins Gtf3c4 and Rab1a at lower values (from 0.05 to 0.250), while other alternatives maintain their position. It is also observed that protein Gtf3c4 is improving its global priority when the weight of position in Ft (solubility alteration) ranking criterion is increasing. These fluctuations are expectable since Gtf3c4 has the first position in PISA assay rank regarding solubility alteration. However, protein Hspb1 maintained the first position at every interval validating the result of the AHP method.

**Table 4.** Fluctuation on global priority of the alternatives when minor variations are done to the criterion position in Ft (solubility alteration) ranking.

| Protein | Global priority fluctuation |  |  |  |  |  |  |  |  |  |  |  |  |  |  |  |  |  |  |
| --- | --- | --- | --- | --- | --- | --- | --- | --- | --- | --- | --- | --- | --- | --- | --- | --- | --- | --- | --- |
|  | 0.050 | 0.100 | 0.150 | 0.200 | 0.250 | 0.300 | 0.350 | 0.400 | 0.450 | 0.500 | 0.550 | 0.600 | 0.650 | 0.700 | 0.750 | 0.800 | 0.850 | 0.900 | 0.950 |
| **Gtf3c4** | 0.190 | 0.208 | 0.226 | 0.244 | 0.262 | 0.280 | 0.298 | 0.316 | 0.334 | 0.352 | 0.370 | 0.388 | 0.406 | 0.424 | 0.442 | 0.460 | 0.478 | 0.496 | 0.514 |
| **Fam98b** | 0.081 | 0.094 | 0.107 | 0.120 | 0.133 | 0.146 | 0.159 | 0.172 | 0.185 | 0.198 | 0.211 | 0.224 | 0.237 | 0.249 | 0.262 | 0.275 | 0.288 | 0.301 | 0.314 |
| **Hspb1** | 0.375 | 0.385 | 0.394 | 0.403 | 0.413 | 0.422 | 0.431 | 0.441 | 0.450 | 0.460 | 0.469 | 0.478 | 0.488 | 0.497 | 0.506 | 0.516 | 0.525 | 0.535 | 0.544 |
| **Rab1a** | 0.236 | 0.242 | 0.249 | 0.256 | 0.263 | 0.270 | 0.276 | 0.283 | 0.290 | 0.297 | 0.303 | 0.310 | 0.317 | 0.324 | 0.330 | 0.337 | 0.344 | 0.351 | 0.357 |
| **Mtpn** | 0.149 | 0.154 | 0.158 | 0.163 | 0.167 | 0.172 | 0.177 | 0.181 | 0.186 | 0.190 | 0.195 | 0.200 | 0.204 | 0.209 | 0.213 | 0.218 | 0.223 | 0.227 | 0.232 |
| **Kiaa0368** | 0.129 | 0.132 | 0.135 | 0.138 | 0.141 | 0.144 | 0.147 | 0.150 | 0.153 | 0.156 | 0.159 | 0.162 | 0.165 | 0.168 | 0.171 | 0.174 | 0.178 | 0.181 | 0.184 |
| **Grhpr** | 0.283 | 0.285 | 0.287 | 0.289 | 0.291 | 0.293 | 0.294 | 0.296 | 0.298 | 0.300 | 0.302 | 0.304 | 0.306 | 0.308 | 0.310 | 0.312 | 0.314 | 0.316 | 0.318 |
| **Cnpy3** | 0.292 | 0.293 | 0.294 | 0.296 | 0.297 | 0.298 | 0.300 | 0.301 | 0.302 | 0.304 | 0.305 | 0.306 | 0.307 | 0.309 | 0.310 | 0.311 | 0.313 | 0.314 | 0.315 |

Moreover, the AHP approach presented here overcomes some limitations from target selections. The parameters included in the semiquantitative analysis are not exclusively based on expert input. The expert knowledge is very valuable, but it is frequently based on reported toxicological studies from only specific targets. For any protein that has not yet studied as possible target, the expert knowledge is limited or absent and the risk of underweighting the relevance of new target is difficult to estimate.

This study offers the opportunity to unravel novel MIEs that could be used to develop AOPs. In the case of TCDD, these new insights are of particular interest since the well-known AOP where it is identified as a stressor, i.e., aryl hydrocarbon receptor activation leading to early life stage mortality, via increased COX-2, has a relevant biological domain of applicability in terms of taxa to all teleost and non-teleost fishes, among other taxa, but it is not applicable to mammals ^32^. Meaning that this AOP cannot be applied to humans.

Although AOPs are not chemical- or stressor-specific, TCDD has been widely used as a compound of reference in toxicology and risk assessment ^33–36^. Therefore, knowing other MIE/AOP that it triggers is of great significance, especially if they can be applicable to humans. This linkage it is not only valuable as an aid for researchers exploring for AOPs that may be relevant to a given stressor but for risk assessment decision-makers evaluating chemicals to enable hazard-based regulation. Additionally, the high throughput unbiased identification of protein targets from all proteins in a studied proteome provides the possibility to fill in missing information of already developed AOPs. Altogether, the integration of AHP approach to support target selection based on PISA target identification could help decision-makers of risk assessment to get access to policy-relevant scientific data and gain in terms of time and resources.

## 4. CONCLUSIONS

We showed that the analysis of chemical-protein interactions by the 2D PISA assay provides an extended list of protein targets, and that the AHP technique improves the process of data curation and target selection with the aim to accurate identify the most appropriate MIE to develop new AOPs. The integration of high-throughput analysis of chemical protein targets by proteomics with target selection by a systematic data curation based on AHP will contribute researchers and regulators to gain time and resources in chemicals assessment.

We expect that this combination of techniques will be a beneficial starting point for the enhancement of the applications of AOPs in chemical risk assessment, such as chemical grouping, chemical prioritization, and integrated approaches to testing and assessment; for the improvement of the development of alternative testing strategies for decision making; and for the advance of the reduction of reliance on animal testing.

## Supporting information

Protein target information used for the pairwise comparison of the alternatives and the computed values from this analysis

## ASSOCIATED CONTENT

### Supporting Information

The Supporting Information is available free of charge.

Protein target information used for the pairwise comparison of the alternatives and the computed values from this analysis (PDF).

## AUTHOR INFORMATION

### Author Contributions

V.L-F. has incorporated additional ideas to the final study, performed the proteomics experiments and the analytical hierarchy process, analyzed the proteomics and nanoDSF data, have written the manuscript and edited the manuscript. A.C.A. has helped with nanoDSF experiment and in the discussion of the results. S.C. has generated the idea, and initial design the study, has supervised the analysis of the results and discussion, have written the manuscript, and was responsible for funding acquisition.

### Notes

The authors declare no competing financial interest.

## ACKNOWLEDGMENTS

This work has been performed with funding from: the ERA-NET Marine Biotechnology project CYANOBESITY that it is cofounding from FORMAS, Sweden grant nr. 2016-02004 (S.C.); the project GOLIATH that has received funding from the European Union’s Horizon 2020 research and innovation programme under grant agreement No 825489 (S.C.); IKERBASQUE, Basque Foundation for Science (S.C.); Basque Government Research Grant IT-971-16 and IT-476-22 (S.C.); Magnus Bergvalls Foundations (S.C.), and the grant for doctoral studies OAICE-75-2017 World Bank counterpart - University of Costa Rica (V.L-F.). All the mass spectrometry analysis has been performed with instrumentation at the LiU MS facility. We thank to Dr. Olatz Fresnedo from University of the Basque Country UPV/EHU, Spain for kindly provide the HepG2 cells.

## ABBREVIATIONS

2D PISA: two dimensions proteome integral solubility alteration
TCDD: 2,3,7,8-tetrachlorodibenzo-p-dioxin
AOP: adverse outcome pathway
MIE: molecular initiating event
MCDM: multi-criteria decision-making analysis
AHP: analytic hierarchy process
DMSO: dimethyl sulfoxide
nLC-MS/MS: nano liquid chromatography-tandem mass spectrometry analysis
FASP: filter aided sample preparation
ACN: acetonitrile
FA: formic acid
Gtf3c4: general transcription factor 3C polypeptide 4
Hspb1: heat shock protein beta-1
Rab1a: ras-related protein Rab-1A
Kiaa0368: proteasome adapter and scaffold protein ECM29
Mtpn: myotrophin
Fam98b: protein FAM98B
Grhpr: glyoxylate reductase/hydroxypyruvate reductase
Cnpy3: protein canopy homolog 3
nanoDSF: nanoscale differential scanning fluorimetry
λ_max_: principal eigenvalue
CI: consistency index
RI: random consistency index
CR: consistency ratio

## FOR TABLE OF CONTENTS ONLY

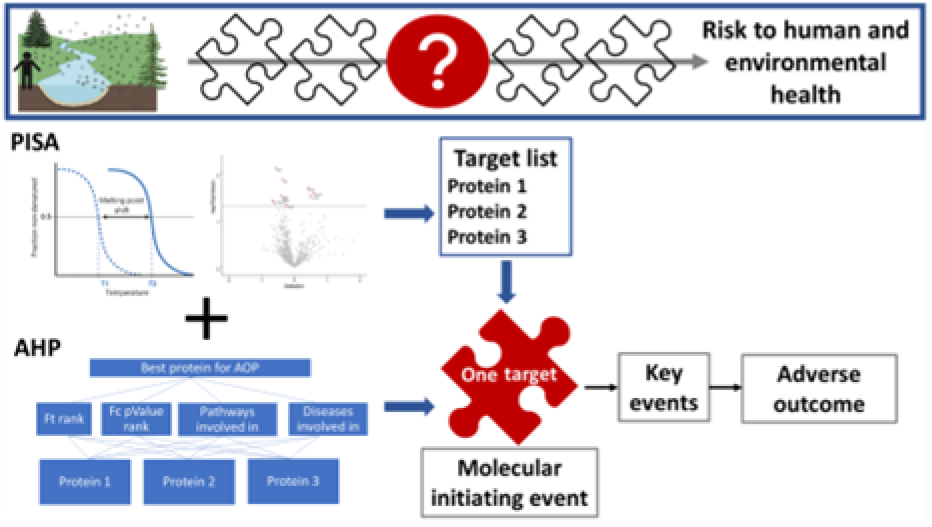

