## Supplementary material for "Proteome integral solubility alteration assay combined with multi-criteria decision-making analysis for developing adverse outcome pathways": Protein target information used for the pairwise comparison of the alternatives and the computed values from this analysis

**Supplementary table 1.** Database containing the information of each protein target (alternatives) for each criterion.

| Protein name | Protein name abbreviation | Position in Ft (solubility alteration) ranking | Position in Fc p-value ranking | Number of diseases where it is involved | Number of reported negative effects on cells/organs when functionality is absent | Relevance of reported negative effects on cells/organs when functionality is absent | References | Number of pathways where it has participation | Relevance of pathways where it has participation | Number of functional and physical protein associations with other protein targets |
|---|---|---|---|---|---|---|---|---|---|---|
| General transcription factor 3C polypeptide 4 | Gtf3c4 | 1 | 1 | 0 | 0 | NA | NA | 1 | Gene expression (Transcription): RNA Polymerase III Transcription | 0 |
| Protein FAM98B | Fam98b | 2 | 7 | 0 | 0 | NA | NA | 1 | Metabolism of RNA: tRNA processing in the nucleus | 0 |
| Heat shock protein beta-1 | Hspb1 | 3 | 4 | 2 | 10 | Charcot-Marie-Tooth axonal neuropathy; distal hereditary motor neuropathy; myopathy; reduced antioxidant capacity; decreased cell viability and increased the incidence of UVB-induced apoptosis; negative effect on cell growth in chinese hamster ovary cells; attenuated contractile force in human bladder smooth muscle cells; impact on calcium homeostasis and muscle energy metabolism; provokes an accumulation of GATA-1 and impairs terminal maturation in CD34+ human cells; ultrastructural abnormalities in the myofibrillar structure in mouse (e.g. destructured myofibrils and higher gaps between myofibrils) | 1-10 | 4 | Signal transduction: signaling by VEGF, extra-nuclear estrogen signaling and MAPK6/MAPK4 signaling; metabolism of ARN: regulation of mRNA stability by proteins that bind AU-rich elements | 0 |
| Ras-related protein Rab-1A | Rab1a | 4 | 5 | 0 | 2 | Synthetic lethal in a haploid human cell line (critical for the survival and/or growth of MDCK cells), Golgi stack and ribbon structures disruption, and endoplasmic reticulum (ER)-to-Golgi trafficking inhibition; perinuclear clustering of early endosomes and delayed transferrin recycling in MDCK cells | 11-12 | 4 | Vesicle-mediated transport: COPI-mediated anterograde transport, COPII-mediated vesicle transport and intra-Golgi and retrograde Golgi-to-ER traffic; metabolism of proteins: RAB geranylgeranylation | 2 |
| Myotrophin | Mtpn | 5 | 2 | 0 | 0 | NA | NA | Protein not found in the sources | NA | 1 |
| Proteasome adapter and scaffold protein ECM29 | Kiaa0368 | 6 | 3 | 0 | 0 | NA | NA | Protein not found in the sources | NA | 1 |
| Glyoxylate reductase/hydroxypyruvate reductase | Grhpr | 7 | 8 | 1 | 1 | Primary hyperoxaluria type II (PH2) | 13 | 1 | Metabolism of aminoacids and derivatives: glyoxylate metabolism and glycine degradation | 0 |
| Protein canopy homolog 3 | Cnpy3 | 8 | 6 | 1 | 1 | Early-onset epileptic encephalopathies, including West syndrome | 14 | 1 | Immune system: trafficking and processing of endosomal TLR | 0 |

**Supplementary table 2.** Pairwise comparison matrices for the alternatives at level 3 of the hierarchy for each criterion and the computed values of local priority vector, principal eigenvalue ($\lambda_{max}$), consistency index (CI), random consistency index (RI), and consistency ratio (CR) for each matrix.

| **Position in Ft (solubility alteration) ranking** | Gtf3c4 | Fam98b | Hspb1 | Rab1a | Mtpn | Kiaa0368 | Grhpr | Cnpy3 | Priority vector |
|---|---|---|---|---|---|---|---|---|---|
| Gtf3c4 | 1 | 2 | 3 | 4 | 5 | 6 | 7 | 8 | 0.360 |
| Fam98b | 1/2 | 1 | 2 | 3 | 4 | 5 | 6 | 7 | 0.259 |
| Hspb1 | 1/3 | 1/2 | 1 | 2 | 3 | 4 | 5 | 6 | 0.188 |
| Rab1a | 1/4 | 1/3 | 1/2 | 1 | 2 | 3 | 4 | 5 | 0.135 |
| Mtpn | 1/5 | 1/4 | 1/3 | 1/2 | 1 | 2 | 3 | 4 | 0.092 |
| Kiaa0368 | 1/6 | 1/5 | 1/4 | 1/3 | 1/2 | 1 | 2 | 3 | 0.061 |
| Grhpr | 1/7 | 1/6 | 1/5 | 1/4 | 1/3 | 1/2 | 1 | 2 | 0.039 |
| Cnpy3 | 1/8 | 1/7 | 1/6 | 1/5 | 1/4 | 1/3 | 1/2 | 1 | 0.026 |

λmax = 9.563 CI = 0.223 RI= 1.41 CR = 0.158

| **Position in Fc p-value ranking** | Gtf3c4 | Fam98b | Hspb1 | Rab1a | Mtpn | Kiaa0368 | Grhpr | Cnpy3 | Priority vector |
|---|---|---|---|---|---|---|---|---|---|
| Gtf3c4 | 1 | 7 | 4 | 5 | 2 | 3 | 8 | 6 | 0.502 |
| Fam98b | 1/7 | 1 | 1/4 | 1/3 | 1/6 | 1/5 | 2 | 1/2 | 0.073 |
| Hspb1 | 1/4 | 4 | 1 | 2 | 1/3 | 1/2 | 5 | 3 | 0.247 |
| Rab1a | 1/5 | 3 | 1/2 | 1 | 1/4 | 1/3 | 4 | 2 | 0.183 |
| Mtpn | 1/2 | 6 | 3 | 4 | 1 | 2 | 7 | 5 | 0.400 |
| Kiaa0368 | 1/3 | 5 | 2 | 3 | 1/2 | 1 | 6 | 4 | 0.318 |
| Grhpr | 1/8 | 1/2 | 1/5 | 1/4 | 1/7 | 1/6 | 1 | 1/3 | 0.041 |
| Cnpy3 | 1/6 | 2 | 1/3 | 2 | 1/5 | 1/4 | 3 | 1 | 0.134 |

λmax = 8.551 CI = 0.079 RI= 1.41 CR = 0.056

| **Number of diseases where it is involved** | Gtf3c4 | Fam98b | Hspb1 | Rab1a | Mtpn | Kiaa0368 | Grhpr | Cnpy3 | Priority vector |
|---|---|---|---|---|---|---|---|---|---|
| Gtf3c4 | 1 | 1 | 1/9 | 1 | 1 | 1 | 1/7 | 1/7 | 0.102 |
| Fam98b | 1 | 1 | 1/9 | 1 | 1 | 1 | 1/7 | 1/7 | 0.102 |
| Hspb1 | 9 | 9 | 1 | 9 | 9 | 9 | 5 | 5 | 0.968 |
| Rab1a | 1 | 1 | 1/9 | 1 | 1 | 1 | 1/7 | 1/7 | 0.102 |
| Mtpn | 1 | 1 | 1/9 | 1 | 1 | 1 | 1/7 | 1/7 | 0.102 |
| Kiaa0368 | 1 | 1 | 1/9 | 1 | 1 | 1 | 1/7 | 1/7 | 0.102 |
| Grhpr | 7 | 7 | 1/5 | 7 | 7 | 7 | 1 | 1 | 0.708 |
| Cnpy3 | 7 | 7 | 1/5 | 7 | 7 | 7 | 1 | 1 | 0.708 |

λmax = 8.042 CI = 0.006 RI= 1.41 CR = 0.004

| **Number of reported negative effects on cells/organs/organisms when functionality is absent** | Gtf3c4 | Fam98b | Hspb1 | Rab1a | Mtpn | Kiaa0368 | Grhpr | Cnpy3 | Priority vector |
|---|---|---|---|---|---|---|---|---|---|
| Gtf3c4 | 1 | 1 | 1/9 | 1/5 | 1 | 1 | 1/3 | 1/3 | 0.100 |
| Fam98b | 1 | 1 | 1/9 | 1/5 | 1 | 1 | 1/3 | 1/3 | 0.100 |
| Hspb1 | 9 | 9 | 1 | 7 | 9 | 9 | 8 | 8 | 0.853 |
| Rab1a | 5 | 5 | 1/7 | 1 | 5 | 5 | 3 | 3 | 0.369 |
| Mtpn | 1 | 1 | 1/9 | 1/5 | 1 | 1 | 1/3 | 1/3 | 0.100 |
| Kiaa0368 | 1 | 1 | 1/9 | 1/5 | 1 | 1 | 1/3 | 1/3 | 0.173 |
| Grhpr | 3 | 3 | 1/8 | 1/3 | 3 | 3 | 1 | 1 | 0.297 |
| Cnpy3 | 3 | 3 | 1/8 | 1/3 | 3 | 3 | 1 | 1 | 0.297 |

λmax = 8.166 CI = 0.024 RI= 1.41 CR = 0.017

| **Relevance of reported negative effects on cells/organs/organisms when functionality is absent** | Gtf3c4 | Fam98b | Hspb1 | Rab1a | Mtpn | Kiaa0368 | Grhpr | Cnpy3 | Priority vector |
|---|---|---|---|---|---|---|---|---|---|
| Gtf3c4 | 1 | 1 | 1/9 | 1/5 | 1 | 1 | 1/9 | 1/9 | 0.097 |
| Fam98b | 1 | 1 | 1/9 | 1/5 | 1 | 1 | 1/9 | 1/9 | 0.097 |
| Hspb1 | 9 | 9 | 1 | 1/7 | 9 | 9 | 1 | 1 | 0.715 |
| Rab1a | 5 | 5 | 7 | 1 | 5 | 5 | 1/7 | 1/7 | 0.374 |
| Mtpn | 1 | 1 | 1/9 | 1/5 | 1 | 1 | 1/9 | 1/9 | 0.097 |
| Kiaa0368 | 1 | 1 | 1/9 | 1/5 | 1 | 1 | 1/9 | 1/9 | 0.097 |
| Grhpr | 9 | 9 | 1 | 7 | 9 | 9 | 1 | 1 | 0.900 |
| Cnpy3 | 9 | 9 | 1 | 7 | 9 | 9 | 1 | 1 | 0.900 |

λmax = 8.496 CI = 0.071 RI= 1.41 CR = 0.050

| **Number of pathways where it has participation** | Gtf3c4 | Fam98b | Hspb1 | Rab1a | Mtpn | Kiaa0368 | Grhpr | Cnpy3 | Priority vector |
|---|---|---|---|---|---|---|---|---|---|
| Gtf3c4 | 1 | 1 | 1/9 | 1/9 | 0 | 0 | 1 | 1 | 0.089 |
| Fam98b | 1 | 1 | 1/9 | 1/9 | 0 | 0 | 1 | 1 | 0.089 |
| Hspb1 | 9 | 9 | 1 | 1 | 0 | 0 | 9 | 9 | 0.801 |
| Rab1a | 9 | 9 | 1 | 1 | 0 | 0 | 9 | 9 | 0.801 |
| Mtpn | 0 | 0 | 0 | 0 | 1 | 0 | 0 | 0 | 0.015 |
| Kiaa0368 | 0 | 0 | 0 | 0 | 0 | 1 | 0 | 0 | 0.005 |
| Grhpr | 1 | 1 | 1/9 | 1/9 | 0 | 0 | 1 | 1 | 0.089 |
| Cnpy3 | 1 | 1 | 1/9 | 1/9 | 0 | 0 | 1 | 1 | 0.089 |

λmax = 8.274 CI = 0.039 RI= 1.41 CR = 0.028

| **Relevance of pathways where it has participation** | Gtf3c4 | Fam98b | Hspb1 | Rab1a | Mtpn | Kiaa0368 | Grhpr | Cnpy3 | Priority vector |
|---|---|---|---|---|---|---|---|---|---|
| Gtf3c4 | 1 | 1/2 | 1/3 | 1/3 | 0 | 0 | 1/5 | 1/3 | 0.062 |
| Fam98b | 2 | 1 | 1/4 | 1/4 | 0 | 0 | 1/5 | 1/4 | 0.116 |
| Hspb1 | 3 | 4 | 1 | 1 | 0 | 0 | 1/3 | 1 | 0.276 |
| Rab1a | 3 | 4 | 1 | 1 | 0 | 0 | 1/3 | 1 | 0.276 |
| Mtpn | 0 | 0 | 0 | 0 | 1 | 0 | 0 | 0 | 0.015 |
| Kiaa0368 | 0 | 0 | 0 | 0 | 0 | 1 | 0 | 0 | 0.005 |
| Grhpr | 5 | 5 | 3 | 3 | 0 | 0 | 1 | 1/3 | 0.416 |
| Cnpy3 | 3 | 4 | 1 | 1 | 0 | 0 | 3 | 1 | 0.306 |

λmax = 8.038 CI = 0.005 RI= 1.41 CR = 0.004

| **Number of functional and physical protein associations with other protein targets** | Gtf3c4 | Fam98b | Hspb1 | Rab1a | Mtpn | Kiaa0368 | Grhpr | Cnpy3 | Priority vector |
|---|---|---|---|---|---|---|---|---|---|
| Gtf3c4 | 1 | 1 | 1 | 1/9 | 1/5 | 1/5 | 1 | 1 | 0.099 |
| Fam98b | 1 | 1 | 1 | 1/9 | 1/5 | 1/5 | 1 | 1 | 0.099 |
| Hspb1 | 1 | 1 | 1 | 1/9 | 1/5 | 1/5 | 1 | 1 | 0.099 |
| Rab1a | 9 | 9 | 9 | 1 | 4 | 4 | 9 | 9 | 0.935 |
| Mtpn | 5 | 5 | 5 | 1/4 | 1 | 1 | 5 | 5 | 0.493 |
| Kiaa0368 | 5 | 5 | 5 | 1/4 | 1 | 1 | 5 | 5 | 0.493 |
| Grhpr | 1 | 1 | 1 | 1/9 | 1/5 | 1/5 | 1 | 1 | 0.099 |
| Cnpy3 | 1 | 1 | 1 | 1/9 | 1/5 | 1/5 | 1 | 1 | 0.099 |

λmax = 8.000 CI = 0.000 RI= 1.41 CR = 0.000
